# Public good diffusion limits microbial mutualism

**DOI:** 10.1101/013144

**Authors:** Rajita Menon, Kirill S. Korolev

## Abstract

Standard game theory cannot describe microbial interactions mediated by diffusible molecules. Nevertheless, we show that one can still model microbial dynamics using game theory with parameters renormalized by diffusion. Contrary to expectations, greater sharing of metabolites reduces the strength of cooperation and leads to species extinction via a nonequilibrium phase transition. We report analytic results for the critical diffusivity and the length scale of species intermixing. We also show that fitness nonlinearities suppress mutualism and favor the species producing slower nutrients.

PACS numbers: 87.23.Cc, 87.23.Kg, 87.15.Zg, 64.60.ah

---

Complex microbial communities are essential for the environment and human health. Microbial functions range from the production of biofuels and the release of powerful greenhouse gasses to the production of cheese and the digestion of food inside our guts. Most of these functions are orchestrated by complex microbial consortia rather than single species [13, 19]. To create and control such multispecies ecosystems, we need to understand the mechanisms that govern microbial coexistence and cooperation.

Heterotrophic cooperation is a common and perhaps the simplest element of complex microbial communities [13–15, 17]. In this two-way cross feeding, each species produces an amino acid or other metabolite necessary for the other species. Heterotrophic cooperation has been previously described by the evolutionary game theory [3, 11], which assumes that microbes interact only with their closest neighbors. However, unlike human societies or bee colonies, microbial communities rarely rely on direct contact. Instead, microbes primarily communicate though diffusible molecules, which rapidly spread in the environment [2, 6, 12, 18, 22]. Because of this diffusive sharing within or between species, such molecules are often termed public goods. The broad understanding of how public good diffusion affects heterotrophic cooperation is still lacking.

Here we explicitly account for the production, consumption, and diffusion of public goods in a model of heterotrophic cooperation. We find that unequal diffusivities of the public goods can significantly favor one of the species and even destroy their cooperation. More importantly, the diffusion of public goods has the opposite effect compared to species migration. Higher migration improves mutualism and stabilizes species coexistence. In contrast, cooperation is lost above a critical diffusivity of public goods, for which we obtain an analytical expression. We also describe the effect of public good diffusion on the spatial distribution of species that is often used to quantify microbial experiments [9, 14, 15, 17]. Our analytical approach is based on computing how public good diffusion renormalizes the strength of selection and thus should be applicable to a variety of more complex models.

Motivated by the experiments on cross-feeding mutualists [14, 15, 17], we consider two species (or strains) A and B producing public goods of type A and B respectively and consuming public goods of the opposite type. These species live in a one-dimensional habitat, which corresponds to the quasi-one-dimensional edge of microbial colonies, where cells actively divide [10]. In simulations, the habitat is an array of islands populated by *N* cells each. This finite carrying capacity sets the magnitude of demographic fluctuations typically termed genetic drift [10]. Nearest-neighbor islands exchange migrants at a rate *m*, which specifies the degree of movement within a microbial colony. In the continuum limit, the evolutionary dynamics of species A and B is described by

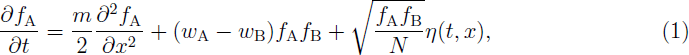

where *t* and *x* are time and position measured in such units that generation time and island spacing are set to 1; *f*_A_(*t*, *x*) and *f*_B_(*t*, *x*) = 1 − *f*_A_(*t*, *x*) are the relative abundances of species *A* and *B*; *w*_*A*_ and *w*_*B*_ are the fitnesses of species *A* and *B* respectively that depend on the local concentration of the public goods; and *η*(*t*, *x*) is a delta-correlated Gaussian white noise. Equation (1) represents the classical stepping-stone model of population genetics [7, 10] and accurately describes population dynamics in microbial colonies [9].

Standard game-theory treatments of microbial interactions assume that the fitnesses *w*_*A*_ and *w*_*B*_ depend on the local abundances of the species rather than the public goods themselves [11]. Here, we relax this restrictive and often unjustified assumption and consider

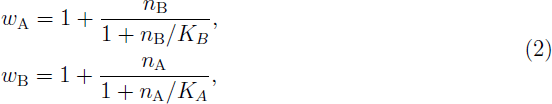

where *n*_A_ and *n*_B_ are the concentrations of the public goods. Although Eq. (2) is known to describe microbial growth well [16], the exact nature of the nonlinear relationship between public good concentrations and fitnesses is of minor importance for our results. Moreover, most aspects of the dynamics can be understood in a much simpler model with *K*_*A*_ = *K*_*B*_ = ∞, where the fitnesses are linear functions of the nutrient concentrations. Note that we chose the units of *n*_*A*_ and *n*_*B*_ to ensure that the numerators in Eq. (2) do not contain any additional constants of proportionality.

In the simplest model, the dynamics of the public goods concentrations are given by the following reaction-diffusion equation:

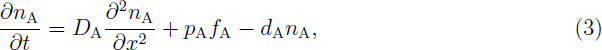

and an analogous equation for *n*_B_. Here, for the public good of type A, *D*_A_ is the diffusivity, *p*_A_ is the production rate, and *d*_A_ is the rate of loss comprised of consumption by both species, spontaneous decay or degradation, and transport outside the region of microbial growth [17]. Both *p*_A_ and *d*_A_ can depend on *n*_A_, *n*_B_ and *f*_A_ in a more realistic model, but numerical simulations suggest that all important aspects of population dynamics are already captured by Eq. (3). Since public good dynamics occurs much faster than cell migration and growth, public good concentrations equilibrate rapidly, i.e. *∂n*_A_/*∂t* ≈ *∂n*_B_/*∂t* ≈ 0. This results in

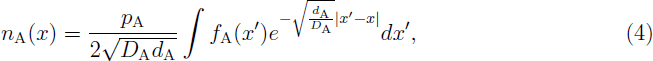

and similarly for *n*_B_. Equation (1-4) have been previously used to simulate cooperatively growing microbial communities [1, 17], but analytical results and broad understanding of the effect of public good diffusion on population dynamics is still lacking.

To understand the overall effect of public good diffusion on microbial mutualism, it is sufficient to consider a simple symmetric case: *p*_A_ = *p*_B_ = *p*, *d*_A_ = *d*_B_ = *d*, *D*_A_ = *D*_B_ = *D*, and *K*_A_ = *K*_B_ = *K* = ∞, which we proceed to analyze by combining Eqs. (1), (2), and (4):

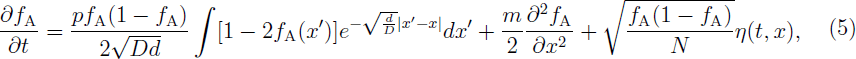

where we eliminated *f*_B_ by using the fact that the relative abundances must sum up to one, i.e. *f*_A_ + *f*_B_ = 1.

For small *D*, the integrand in the first term of Eq. (5) is peaked around *x'* = *x*, so one can simplify the equation by expanding *f*_A_(*x'*) in Taylor series around *x*. To the first order, the selection term becomes *sf*_A_(1 − *f*_A_)(1/2 − *f*_A_), where *s* = 2*p*/*d* is the selection coefficient. Thus, when diffusion is very slow, our model of mutualism reduces to the standard game theory formulation with frequency-dependent selection. Population dynamics in this limit are controlled by a dimensionless quantity *S* = *smN*^2^, which we refer to as the strength of the mutualism. When *S* exceeds a critical value of order one *S*_c_, the mutualism is stable, and the two species coexist [11]. In contrast, when *S* < *S*_c_, selection for coexistence is not strong enough to overcome local species extinctions due to genetic drift, and the population becomes partitioned into domains exclusively occupied by one of the two species (see inset in Fig. 1). Generically, this demixing phase transition belongs to the universality class of directed percolation (DP), but the special symmetric case, when all model parameters are the same for the two species, is in the DP2 or generalized voter universality class [11].

**FIG. 1:**
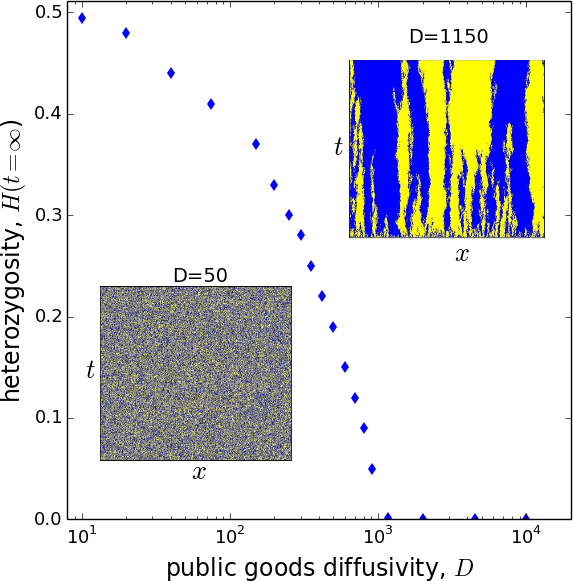
Mutualism is destroyed by genetic drift at high nutrient diffusivities. Equilibrium values of local heterozygosity decay as the diffusivity of the metabolites increases, and the species coexistence is completely lost above a critical diffusivity *D*_c_. The two insets show how the distribution of species labeled by different colors varies with position (x-axis) and time (y-axis) for small diffusivities (left) and just below the critical diffusivity (right). One can clearly see the mixed and demixed states observed experimentally [14, 15, 17]. Here *N* = 200, *m* = 0.1, *p* = 0.001, *d* = 1, and *K* = 1. The habitat consisted of 10^4^ islands and was observed after 10^6^ generations starting from a well-mixed state.

Expansion to the next order of the integral in Eq. (5) contributes an additional term −2*pDd*^−2^ *f*_A_(1 − *f*_A_)*∂*^2^ *f*_A_/*∂x*^2^, which effectively reduces migration *m* and, therefore, the strength of mutualism *S*. To test whether public good diffusion weakens mutualism, we quantified species coexistence by the average local heterozygosity *H*(*t*) = 〈2*f*_A_(*t*, *x*)*f*_B_(*t*, *x*)〉, which equals 1/2 for strongly intermixed species and 0 for species that are spatially segregated. In computer simulations, we then observed that equilibrium values *H* indeed decrease with *D*, and mutualism is lost for diffusivities above a certain value *D*_c_ (Fig. 1). Note that the loss of mutualism in our model is due to genetic drift rather than the proliferation of nonproducers (or cheaters), which are commonly considered in the context of public goods[1].

The small *D* expansion is only valid when *f*_A_ is slowly varying in space, but microbial communities are often found at low values of spatial intermixing when individual species appear as clusters or domains [8, 14, 15, 17, 21]. Indeed, Fig. 1 shows that intermediate values of local heterozygosity are observed as *D* changes by almost two orders of magnitude. To understand population dynamics in this important regime, we consider large diffusivities when the size of the domain boundaries *L*_b_ is much smaller than the nutrient length scale 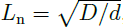 i.e. the typical distance between the locations of nutrient production and consumption. To the first order, the size of the domain boundaries is the same as in the neutral model with no mutualism, for which *L*_b_ = *mN* [4]. We will also assume that the population is sufficiently close to the demixing phase transition so that the distance between domain boundaries *L*_d_ is much greater than *L*_n_, as it is commonly observed experimentally [14, 15, 17].

When *L*_d_ ≫ *L*_n_ ≫ *L*_b_, one can solve Eq. (5) near the domain boundary located at *x* = 0 by assuming that *f*_A_(*x*) is a step function, i.e. *f*_A_(*x*) = 1 for *x* < 0 and *f*_A_(*x*) = 0 for *x* > 0. The solution yields

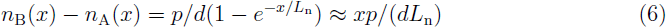

for *x* << *L*_n_.

Since the selection term in Eq. (5) vanishes when *f*_A_(1 − *f*_A_) = 0, only the fitness differences at the domain boundary (when *x* ≈ *L*_b_) affect population dynamics. Near the boundary *f*_A_(*x*) ≈ 1/2 − *x*/*L*_b_. Hence, by eliminating *x* from Eq. (6), we can again recast the selection term in the form *s*_eff_*f*_A_(1 − *f*_A_)(1/2 − *f*_A_), where the effective strength of selection *s*_eff_ ∼ (*p*/*d*)(*L*_b_/*L*_n_) is reduced by a factor of 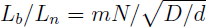 compared to the model with *D* = 0.

Our finding that higher diffusivities of the public goods reduce the effective strength of selection explains the decrease of *H* with *D* in Fig. 1 and provides a way to estimate the critical diffusivity *D*_c_ above which mutualism is lost. Indeed, when *D* = *D*_c_, we expect that the strength of mutualism *S* approaches its critical value as well. Thus, 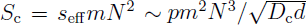, and *D*_c_ ∼ *s*^2^*m*^4^*N*^6^*d*^3^, where *s* = 2*p*/*d* is the strength of selection in the model without public good diffusion. Surprisingly, we find that population density and migration have a much stronger effect on the critical nutrient diffusivity than natural selection. Our simulation results are in excellent agreement with these predictions; see Fig. 2.

**FIG. 2:**
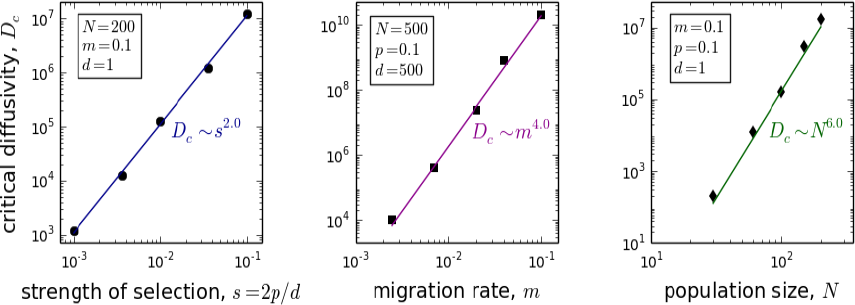
Simulations (points) confirm our analytical predictions (lines) that critical diffusivity increases as a square of the selection strength *s*, fourth power of migration rate *m*, and sixth power of the population size *N*. All data are from simulations on a lattice of 10^4^ islands over 10^6^ generations with *K* = 1.

Since the model with *D* > 0 is equivalent to that with *D* = 0 provided the strength of selection *s* is renormalized, many of the results from the evolutionary game theory can be generalized for microbial communities with diffusible public goods. The size of the domains formed by the species *L*_d_ is of particular interest because it is used in the experiments to quantify the degree to which the two species benefit from their mutualistic interactions. Previous studies suggested that *L*_d_ ∼ *D*^1/4^ [17] or *L*_d_ ∼ *D*^1/5^ [14]; however, we find that such scalings are unlikely because *L*_d_ becomes large only close to the underlying phase transition, where *L*_d_ ∼ (*D*_c_ − *D*)^*ν*⊥^. The exponent *ν*_⊥_ is that of a spatial correlation length and is determined by the universality class of the phase transition. In our model, any species asymmetry results in DP universality class and *ν*_⊥_ = 1.096854(4), while, when all model parameters are the same for the two species, the dynamics is in DP2 universality class and *ν*_⊥_ = 1.83(3) [5, 20]. Simulations confirm this expectation as shown in Fig. 3.

**FIG. 3:**
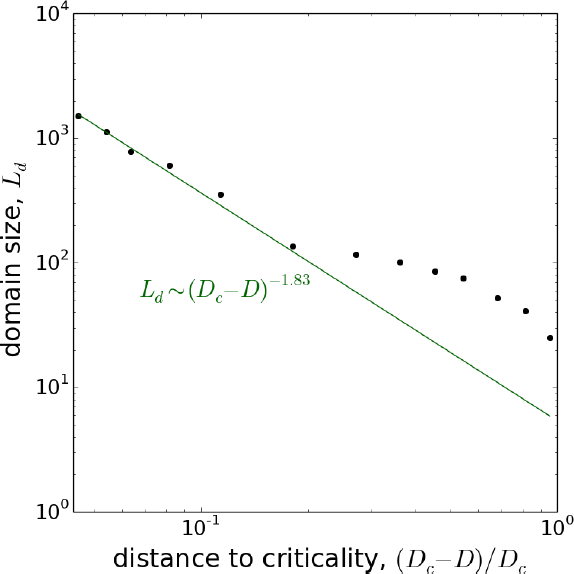
The scale of species intermixing is controlled by the underlying nonequilibrium phase transition. The dots are the simulation data, and the line is the fit of the expectation that *Ld* ∼ (*D*_c_ − *D*)^*v*⊥^, where *v*_⊥_ = 1.83 is the correlation-length exponent for the generalized voter (or directed percolation 2) critical point [5, 20]. The simulations were carried out on a lattice of 10^4^ islands for 10^6^ generations. Here, *N* = 200, *m* = 0.1, *p* = 0.001, *d* = 1, and *K* = 1.

Next we turn to the effects of species asymmetries on the population dynamics. When *D* = 0 and the model reduces to that of frequency-dependent selection, the asymmetries in nutrient production and decay rates result in the selection term of the form *sf*_A_(1 − *f*_A_)(*f*\* − *f*_A_), where the preferred fraction *f*\* no longer equals 1/2. Population dynamics in this limit have been previously analyzed in Ref. [11] with the main conclusion that species asymmetry substantially weakens mutualism and species A is favored if *f*\* > 1/2 while species B is favored otherwise. We find that *D* > 0 does not alter these results.

The asymmetry in the public good diffusivities is more subtle. Upon repeating the steps leading to Eq. (5) but for *D*_A_ ≠ *D*_B_ one still finds that the resulting equation is invariant under the exchange of species labels, i.e. *f*_A_ → 1 − *f*_A_. Hence, if only the public good diffusivities are different between the species, then none of the species is expected to dominate the other. This conclusion however holds only when the fitnesses are linear functions of the nutrient concentrations, i.e. *K* = ∞. For lower values of *K*, we find that the species producing public goods that diffuse more slowly dominates the other species (Fig. 4). As *K* is decreased further, the population undergoes the demixing phase transition described above, and one of the species becomes extinct.

The effects of fitness nonlinearities and public good diffusivities can be easily understood by considering the population dynamics close to the domain boundary. The dominant species is determined by whether species A is more likely to invade the space occupied by species B or species B is more likely to invade the space occupied by species A. To make the argument more clear let us assume that *D*_A_ = 0 and *D*_B_ = ∞, then the concentration of public good B is the same everywhere while the concentration of public good A is high inside the domain comprised of species A and zero outside. As a result, the fitness of species A is the same everywhere, while the fitness of species B is low in its own domain and high in the domain occupied by species A. The nonlinearity in Eq. (2) makes fitness changes at low nutrient concentrations much more pronounced than at high nutrient concentrations. Thus, the advantage that the species B has over A in the domain occupied by species A (where *n*_A_ is high) is smaller than the advantage that species A has over B in the domain occupied by species B (where *n*_A_ = 0). In consequence, species A with lower public good diffusivity should dominate species B in agreement with the simulations (Fig. 4).

**FIG. 4:**
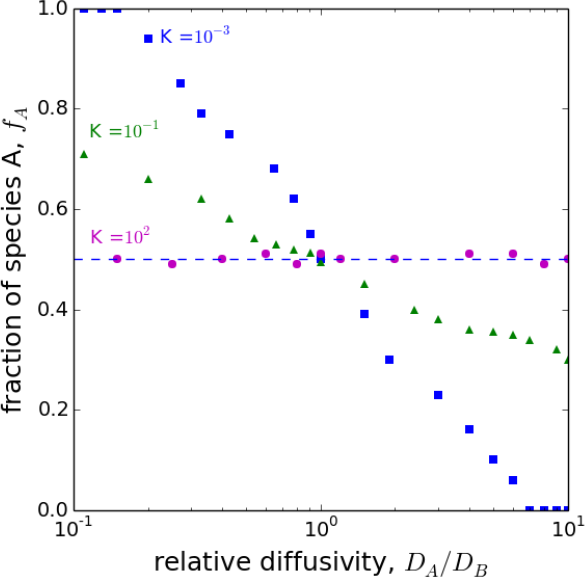
Nonlinearity due to the diminishing effect of public goods on fitness benefits the species producing public goods with smaller diffusivity. The degree of fitness nonlinearity decreases as a function of parameter *K* in Eq. (2). When *K* is very large, the model is linear and the differences in metabolite diffusivities have no affect on the relative species abundances (magenta dots). As the fitness nonlinearity is increased, the species with the lower diffusivity dominates (green triangles). This trend continues as *K* is decreased further, but, in addition, species coexistence and mutualism are lost when the metabolite diffusivities become too unequal (blue squares). These simulations were run for 10^5^ generations on a lattice of 500 islands; *N* = 200, *m* = 0.1, *p* = 0.001, and *d* = 1.

In summary, we demonstrated that the main effect of public good diffusion is the reduction of the effective strength of natural selection, which can lead to the loss of mutualism via a nonequilibrium phase transition. The distance to this phase transition controls the size of the domains formed by the species, a quantity of prime interest in empirical studies. In addition, differences in the diffusivities of the public goods could have a profound effect on the population dynamics. The effect of these differences depends on the fitness and other nonlinearities and results in the selective advantage for one of the species. Our work provides a theory for the phenomena observed in recent experimental studies [14, 15, 17] and could potentially explain why cooperatively growing microbes modulate the diffusivities of their public goods [12].

